# White matter tract microstructure, macrostructure, and associated cortical gray matter morphology across the lifespan

**DOI:** 10.1101/2023.09.25.559330

**Authors:** Kurt G Schilling, Jordan A. Chad, Maxime Chamberland, Victor Nozais, Francois Rheault, Derek Archer, Muwei Li, Yurui Gao, Leon Cai, Flavio Del’Acqua, Allen Newton, Daniel Moyer, John C. Gore, Catherine Lebel, Bennett A Landman

## Abstract

Characterizing how, when and where the human brain changes across the lifespan is fundamental to our understanding of developmental processes of childhood and adolescence, degenerative processes of aging, and divergence from normal patterns in disease and disorders. We aimed to provide detailed descriptions of white matter pathways across the lifespan by thoroughly characterizing white matter *microstructure*, white matter *macrostructure*, and morphology of the *cortex* associated with white matter pathways. We analyzed 4 large, high-quality, publicly-available datasets comprising 2789 total imaging sessions, and participants ranging from 0 to 100 years old, using advanced tractography and diffusion modeling. We first find that all microstructural, macrostructural, and cortical features of white matter bundles show unique lifespan trajectories, with rates and timing of development and degradation that vary across pathways – describing differences between types of pathways and locations in the brain, and developmental milestones of maturation of each feature. Second, we show cross-sectional relationships between different features that may help elucidate biological changes occurring during different stages of the lifespan. Third, we show unique trajectories of age-associations across features. Finally, we find that age associations during development are strongly related to those during aging. Overall, this study reports normative data for several features of white matter pathways of the human brain that will be useful for studying normal and abnormal white matter development and degeneration.

## Introduction

Human brain white matter is composed of fiber pathways that connect different components of the neural system. Because fiber pathways determine the brain’s functional organization, they play a critical role in nearly every aspect of cognition and behavior. These white matter tracts undergo significant changes throughout the lifespan, and characterizing how, when and where these changes occur is critical to our understanding of developmental processes of childhood and adolescence, degenerative processes of aging, and the divergence from normal patterns of change in disease and disorder.

### White Matter Microstructure

Diffusion tensor imaging (DTI) [1, 2] shows robust increases in fractional anisotropy (FA) and decreases in axial, radial, and mean diffusivities (AD, RD, MD) in childhood and adolescence [3-5], with opposite trends in healthy aging [6, 7]. Lifespan studies reveal complex nonlinear patterns, where white matter microstructure typically reaches maturation between the early twenties to late thirties [8-10]. Multi-compartment models of diffusion may offer increased biological specificity. One such model, the Neurite Orientation Dispersion and Density Imaging (NODDI) [11] model provides estimates of neurite density (intracellular volume fraction; ICVF), extracellular water diffusion (isotropic volume fraction; ISOVF), and tract orientation dispersion (OD). NODDI has been used to probe biological changes in infancy [12], childhood [13], and aging [14], but lifespan studies [15] have been reported less due to more advanced MRI acquisition requirements and limited sample sizes.

### White Matter Macrostructure

Beyond the microstructural measures provided by diffusion MRI, the *macrostructural* features of white matter pathways may play a pivotal role along the aging continuum. For example, studies suggest large initial increases in white matter pathway volume followed by a plateau and decreases at later ages, with rates and timing of development and degradation varying across pathways, with strong relationships between white matter macrostructure and the underlying microstructural measures [8]. Moreover, other macrostructural features – including additional measures of tract volumes, areas, and lengths - have been recently measured [16], but not yet characterized across the lifespan.

### Cortical Gray Matter Morphometry

Cortical thickness, volume, and surface area have been well-characterized [17], showing developmental and aging patterns that vary regionally in the brain [18-22], associations with behavior and cognition [23, 24], and differences in neuropsychiatric disorders [25, 26]. However, cortical characterizations are typically performed independently of the underlying white matter connections, and the complex relationships between white matter pathways of the brain and their associated cortical structure has been less thoroughly investigated [27-29].

### A full characterization of white matter pathways across the lifespan

Previous studies have often been limited by sample size, age range, and number of features, or did not investigate specific white matter pathways of the human brain. Because of this, few studies have investigated the links between the developmental processes of childhood and later degenerative processes of aging [9, 30, 31]. In the current study, we analyze 2789 imaging datasets on participants ranging from 0 to 100 years old to segment and quantify microstructure, macrostructure, and cortical features of 63 white matter pathways. Specifically, we address four questions: (1) How are these brain features associated with age throughout the lifespan? (2) How are different features related, and how do these relationships vary across the lifespan? (3) Are age associations of different features related to each other? and (4) Do rates of white matter pathway development influence rates of brain degeneration later in life? In particular, we investigate a theory of retrogenesis known as the “gain predicts loss” hypothesis that proposes that pathways with features that develop faster correspond to those that also decline faster in aging [9, 30, 31].

## Methods

### Datasets

The data used in this study come from the Human Connectome Project (Essen et al. 2012), which aims to map the structural connections and circuits of the brain and their relationships to behavior by acquiring high-quality magnetic resonance images. We used diffusion MRI data from the Baby Connectome Project [32], the Human Connectome Project Development (HCP-D) study, the Human Connectome Project Young Adult (HCP-YA) study, and the Human Connectome Project Aging (HCP-A) study. Within this manuscript, we will refer to these as (capitalized) *Infant*, *Development*, *Young Adult*, and *Aging* cohorts, respectively.

The Infant cohort was composed of 259 participants and 543 imaging sessions (1-5 sessions per participant) aged between 0 and 5 years. The Development cohort was composed of 652 participants aged 5 to 21 years. The Young Adult cohort was composed of 1206 participants aged 21 to 35 years. The Aging cohort was composed of 722 participants aged 35 to 100 years. After quality assurance (removal of datasets with excessive diffusion artifacts, failure of tractography, and/or failure of FreeSurfer [33] or Infant FreeSurfer [34] on the structural images), the pooled dataset was composed of 2789 imaging sessions and spanned the ages of 1 week to 100 years old. Data are summarized in **Table 1**.

**Table 1.**
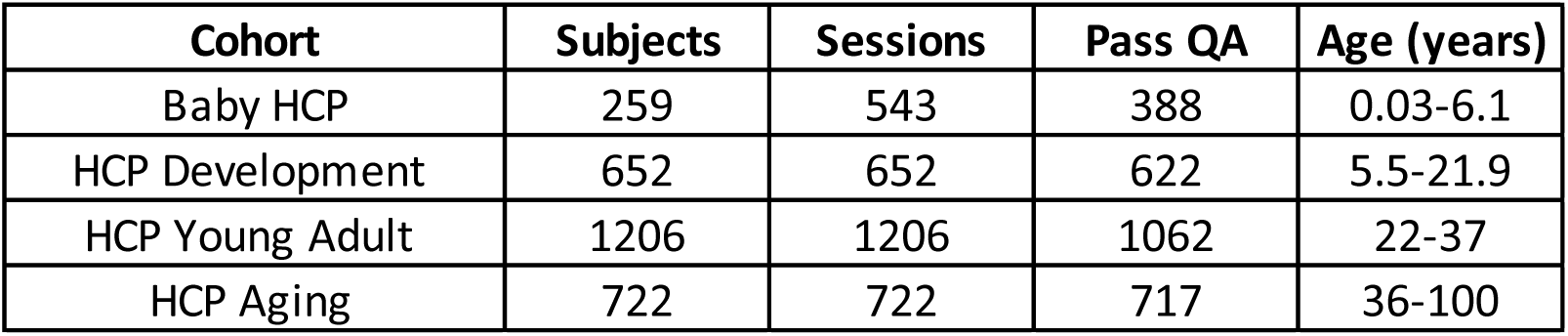
*Data used in this study come from the Human Connectome Project (Essen et al. 2012) initiatives—the Baby Connectome Project (Infant), the Human Connectome Project Development (Development) study, the Human Connectome Project Young Adult (Young Adult) study, and the Human Connectome Project Aging (Aging) study. QA: Quality Assurance check for data quality + successful structural and diffusion processes. Note Baby HCP Cohort is a longitudinal dataset*.

| Cohort | Subjects | Sessions | Pass QA | Age (years) |
| --- | --- | --- | --- | --- |
| Baby HCP | 259 | 543 | 388 | 0.03-6.1 |
| HCP Development | 652 | 652 | 622 | 5.5-21.9 |
| HCP Young Adult | 1206 | 1206 | 1062 | 22-37 |
| HCP Aging | 722 | 722 | 717 | 36-100 |

The diffusion MRI acquisitions were slightly different for each dataset and tailored towards the population under investigation. For Development and Aging cohorts, a multi-shell diffusion scheme was used, with b-values of 1500 and 3000 s/mm^2^, sampled with 93 and 92 directions, respectively (24 b=0). The in-plane resolution was 1.5 mm, with a slice thickness of 1.5 mm. For the young adult cohort, the minimally preprocessed data [35] (Glasser et al. 2013) from Human Connectome Projects (Q1-Q4 release, 2015) were acquired at Washington University in Saint Louis and the University of Minnesota [36] (Essen et al. 2012) using a multi-shell diffusion scheme, with b-values of 1000, 2000, and 3000 s/mm^2^, sampled with 90 directions each (18 b=0). The in-plane resolution was 1.25 mm, with a slice thickness of 1.25 mm. The Infant cohort typically used a 6-shell sampling scheme with b-values of 500, 1000, 1500, 2000, 2500, and 3000 s/mm^2^, sampled with 9, 12, 17, 24, 34, and 48 directions, respectively (14 b=0). Depending on compliance, however, a protocol matched to the Development cohort was sometimes also used for the Infant cohort (see [32] for discussion on acquisition). The in-plane resolution was 1.5 mm, with a slice thickness of 1.5 mm.

For all diffusion data, susceptibility, motion, and eddy current corrections were performed using TOPUP and EDDY algorithms from the FSL package following the minimally preprocessed HCP pipeline [35].

Structural images with T1-weighting for all cohorts were acquired with an MPRAGE sequence, with a resolution of 0.8 mm isotropic for the Infant, Development, and Aging cohorts, and resolution of 0.7 mm isotropic for the Young Adult cohort.

### Tractography and Tract Features

For every session, sets of white matter pathways were virtually dissected using the TractSeg [37] automatic white matter bundle segmentation algorithm. TractSeg was based on convolutional neural networks and performed bundle-specific tractography based on a field of estimated fiber orientations [37]. From the TractSeg outputs, we selected 63 white matter bundles for analysis, including association, commissural, thalamic, striatal, and projection and cerebellar pathways. A list of pathways and acronyms is given in the appendix.

Features of microstructure, macrostructure, and connecting cortical features were extracted for each pathway (**Figure 1**). For microstructure, we first fit the DTI model to all participants using the FSL software *dtifit* algorithm (limiting fitting to b<=1500 s/mm^2^ only), resulting in voxel-wise maps of FA, MD, AD, and RD. We then fit the NODDI model [11] using the scilpy tractography toolbox (https://github.com/scilus/scilpy), resulting in voxel-wise maps of ICVF, ISOVF, and OD. For each tract, and each participant, these values were simply averaged across all voxels in the entire tract.

**Figure 1.**
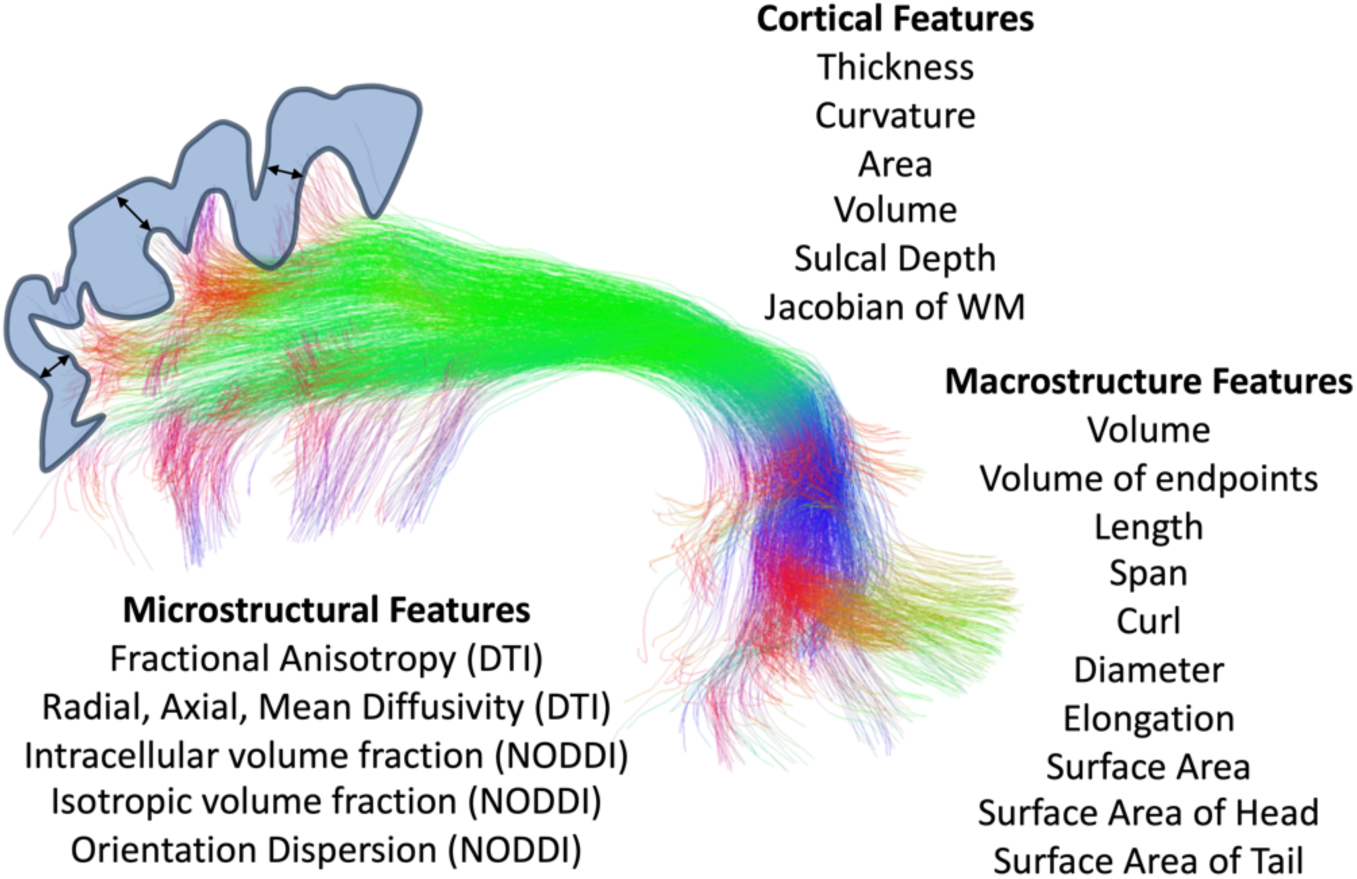
*Microstructural, macrostructural, and cortical features associated with each of 63 white matter bundles*.

For macrostructure, we used the *scil_evaluate_bundles_individual_measures* script from the scilpy toolbox to derive 10 shape features of each tract. Briefly, these features are inspired by [16], and include: volume (total tract volume, mm3); volume of endpoints (volume of voxels containing bundle endpoints, mm3); length (length of bundle, mm); span (distance between two ends of the bundle, mm); curl (measure of curvature ranging from 1 to infinity); diameter (bundle diameter when approximated as a cylinder); elongation (ratio of length to diameter); surface area (total surface area of bundle); surface area of head (surface area of beginning of bundle, i.e., the end that is most left/posterior/inferior); surface area of tail (surface area of end of bundle, i.e., the end that is most right/anterior/superior).

Finally, cortical features associated with each bundle were computed by probing the endpoints of each streamline in a bundle (i.e., in the cortex or at the white/gray matter interface), and averaging these values across all streamlines. Cortical features were calculated from FreeSurfer run on the development, young adult, and aging cohorts, and Infant FreeSurfer run on the Infant cohort. FreeSurfer resulted in surface-based and voxel-based measures along the cortical ribbon of six cortical features: cortical thickness, volume, area, curvature (sulci have positive curvature, gyri negative, with sharper curvature indicated by higher absolute value), Jacobian of white matter (which computes how much the white matter surface needs to be distorted to register to the spherical atlas), and sulcal depth (distance in mm from mid-surface between gyri/sulci, sulci have positive values, gyri have negative values). We note that Infant FreeSurfer did not output volume nor Jacobian of white matter values. For each bundle, we used the *tckmap* feature of the MRTrix3 software package to probe the cortical thickness at the endpoints of every streamline, averaging each feature at both ends (for example averaging cortical thickness at the beginning and end of each streamline), and taking the average of all streamlines. This gives us features such as ‘the cortical thickness associated with the left Arcuate Fasciculus’, for example.

### Associations with Age

To investigate how brain features are associated with age throughout the lifespan we quantified the slopes of age-associations in infancy, development, young adult, and aging cohorts. For every feature of every bundle, we had data points from 2789 imaging sessions with participants ranging in age from 0 to 100 years old. When visualizing the raw data plotted against age, we noticed that the lifespan plots did not result in a smooth trajectory with age due to an offset (in nearly every feature) in the young adult cohort (which had different image resolution and diffusion acquisition parameters than the other three cohorts). Traditionally, data acquired from different sites and acquisitions would be harmonized, for example using the ComBat adjustment method [38] to reduce scanner and acquisition effects. However, there is no overlap in the covariate (age) between cohorts, making this, and other, harmonization methods unfeasible. Thus, we applied a simple statistical adjustment based on continuity assumptions to harmonize data across cohorts. See Supplementary Documentation and **Supplementary Figure 1** for a detailed description of this methodology.

Linear and quadratic fits are common in studies of development and aging, with quadratic fits common in lifespan studies due to the expected reversal and U-shaped curve of most features. However, quadratic fits may not be ideal because they restrict slopes on either side of the peak/minimum to be the same. Thus, less restrictive Poisson curve fits have become popular, especially in the diffusion MRI literature. Despite this, we found that Poisson fits still did not fit the highly nonlinear trends well, particularly during infancy. For this reason, all features across age were analyzed using covariate-adjusted restricted cubic spline regression (C-RCS) [39], a flexible approach to model nonlinear relationships between variables. Here, we used knots at 2, 4, 22, 35, 75, and 90 years of age, based on expected developmental shifts in volumetry [40], with five to six knots being common as a compromise between flexibility and overfitting [41]. The 95% confidence intervals of volumetric trajectories of each tissue/region of interest are derived by deploying C-RCS regression on 10,000 bootstrap samples.

From these C-RCS curves, the Difference per Year and Percent Difference per Year were calculated as measures of cross-sectional associations with age (i.e., cross sectional change per year or cross-sectional percent change per year). The peak/minimum of each curve can be derived, as well as the 95% confidence intervals of each peak. Results in this manuscript are displayed as curves across the lifespan and also summarized by binning each age-group pre-determined based on those in the HCP cohorts, infant (0-5), development (5-21), young adult (22-35), aging (36+).

### Relationships between features across a population

To examine the cross-sectional relationships between microstructure/macrostructure and cortical features, for each pathway we calculated the correlation coefficient between each feature and all other features (i.e., the correlation coefficient between 2789×1 vector of feature 1 and the 2789×1 vector of feature 2). We additionally performed partial linear correlation between all features while controlling for subject age and sex. To account for multiple comparisons, all statistical tests incorporated a false discovery rate at 0.05 to determine statistically significant relationships between features.

### Relationships between age-associations in features

We additionally aimed to ask two questions probing the relationships between *age associations* in features. First, we asked whether greater/lesser age-associations in one feature correspond to greater/lesser age-associations in another feature. Within each cohort, for all pathways we calculated the correlation coefficients to relate the difference per year of each feature to all other features (i.e., the correlation coefficient between the 63×1 vector of difference per year of feature 1 for all pathways and the 63×1 vector of difference per year of feature 2 for all pathways).

Second, we asked whether features that develop faster correspond to those that decline faster/slower in aging. To do this, for a given feature, we calculated the correlation coefficients to relate the cross-sectional change per year of each feature in the development cohort to the cross-sectional change per year of the same feature in the aging cohort (i.e., the correlation coefficient between the 63×1 vector of difference per year of feature 1 for all pathways in the development cohort and the 63×1 vector of difference per year of the same feature 1 for all pathways in the aging cohort). Again, statistical tests incorporated a false discovery rate at 0.05.

## Results

### How are white matter features associated with age throughout the lifespan?

#### Microstructure

Example lifespan trajectories of FA and ICVF are shown in **Figure 2**, along with bundles visualized and colored based on the % difference per year for each cohort. Age related trends are nonlinear, yet smoothly varying, across the lifespan. In both cases, large cross-sectional increases in both measures are observed during infancy, continuing throughout childhood and adolescence, leveling off in young adulthood, and (typically) decreasing in aging. Patterns of change also vary according to pathway location and type, with an anterior-to-posterior gradient of both measures during infancy, and also differences in the low negative age associations of FA in association and commissural pathways in young adulthood, with opposite trends in most thalamic, striatal, and projection pathways. Similar spatial patterns are observed for both FA and ICVF. Trajectories of additional microstructural features are given as **Supplementary Figure 2**.

**Figure 2.**
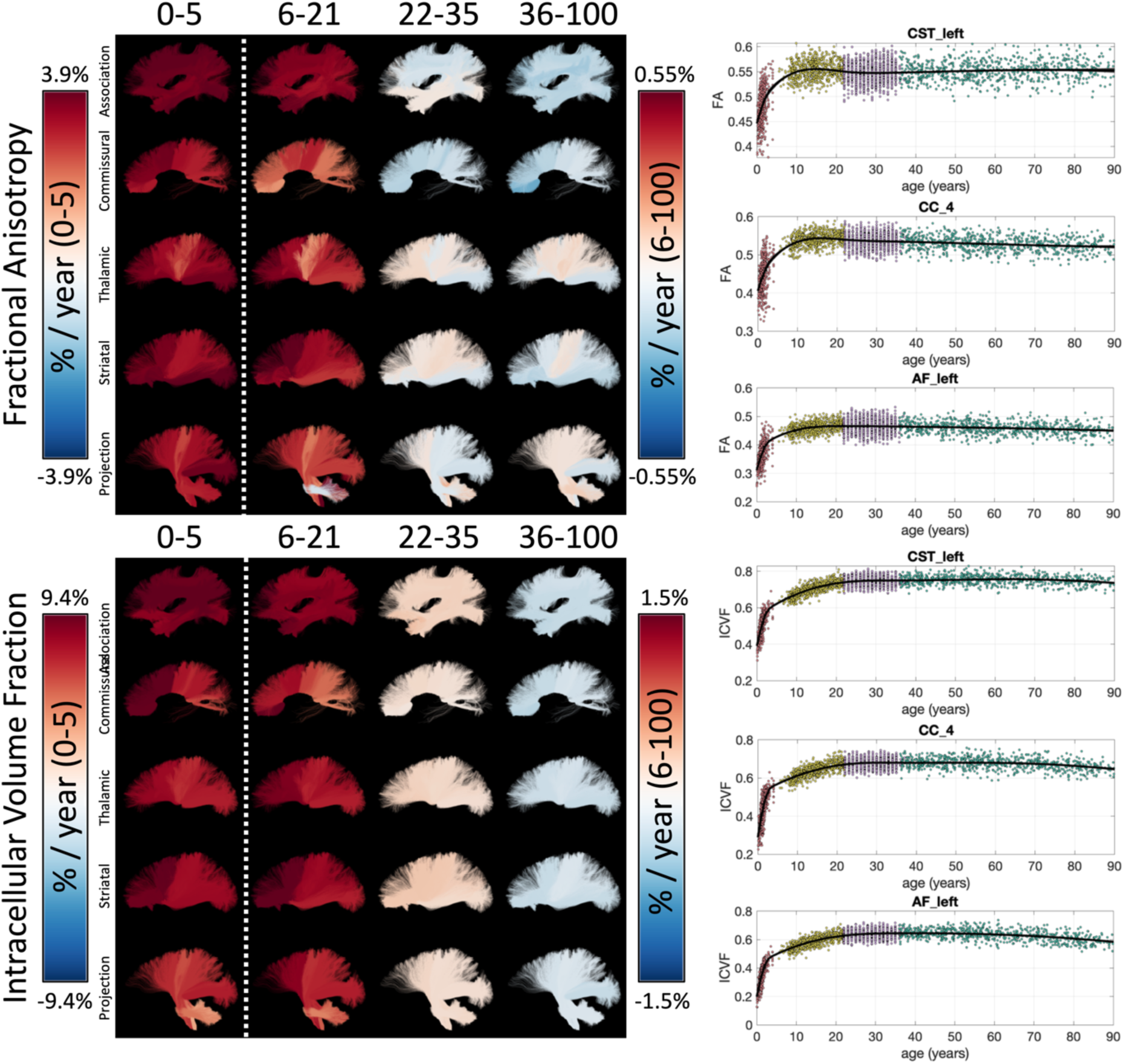
*White matter microstructure shows unique lifespan trajectories. Lifespan trajectories of FA (top) and ICVF (bottom) are shown colored based on the % change per year for each cohort (left) and data points are plotted against age for three selected pathways (right). Note 0- 5 cohort is displayed on a different color scale than other cohorts*.

**Figure 3** summarizes the % difference per year of all white matter microstructure features, for all bundles, across all four cohorts. Here, each cohort is visualized on a different scale to highlight trends across features and pathways. In both Infant and Development cohorts there are large cross-sectional increases in anisotropy, decreases in diffusivities (MD, AD, RD), increases in ICVF, and decreased dispersion. Trends are similar between these two cohorts, age associations of Development are ∼5x smaller in magnitude. Negative age associations of diffusivities continue into Young Adulthood, along with most pathways showing continued positive age associations in ICVF. However, OD now has positive age associations, while FA shows heterogenous associations across pathways, in agreement with visualizations in **Figure 2**. Finally, the Aging cohort displays strong positive associations with age for diffusivities, and negative associations with age in anisotropy, ICVF, and OD. Notably, Thalamic and Projection pathways tend to display opposite patterns with trends opposite to most association pathways for microstructure features of orientation and complexity (FA, OD) but similar trends with features of size and diffusivities (MD, AD, RD, ICVF, ISOVF).

**Figure 3.**
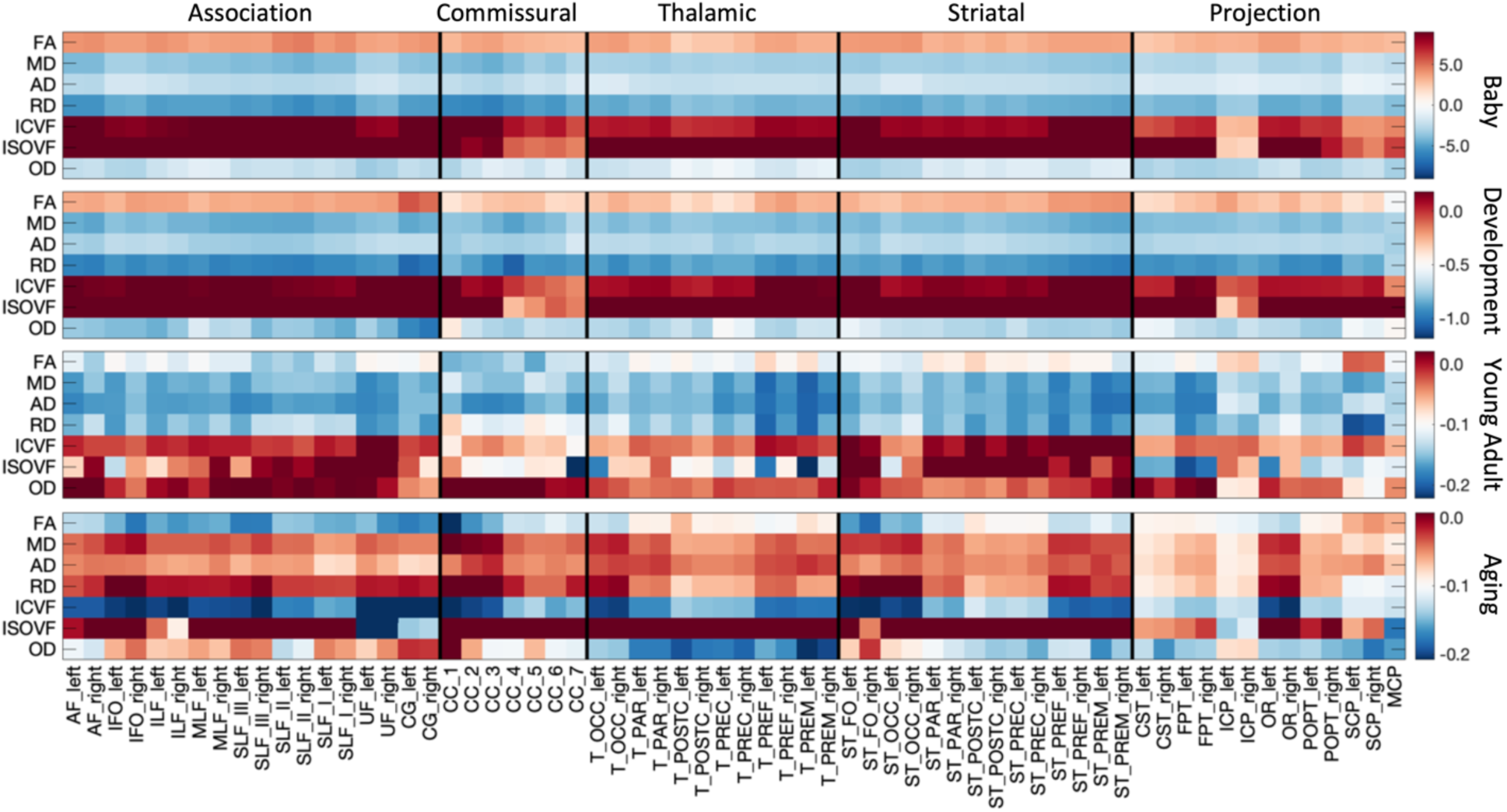
*White matter microstructure shows different associations with age across pathways and across different lifespan stages. The % change per year of all white matter microstructure features, for all bundles is shown for each of the four cohorts. Note, each cohort is visualized on a different scale to highlight trends across features*.

#### Macrostructure

Example lifespan trajectories for macrostructural features of bundle volume and bundle diameter are shown in **Figure 4**, again, with bundles colored based on the % difference per year. Intuitively, total volume and diameter show very similar trends across the lifespan. Again, large cross-sectional increases in volume occur during infancy, and continue into childhood/adolescence for most pathways, although at an order of magnitude lower % difference per year. Both features begin decreasing in Young Adulthood, with trends continuing into Aging, with many bundles experiencing accelerated negative associations at older ages.

**Figure 4.**
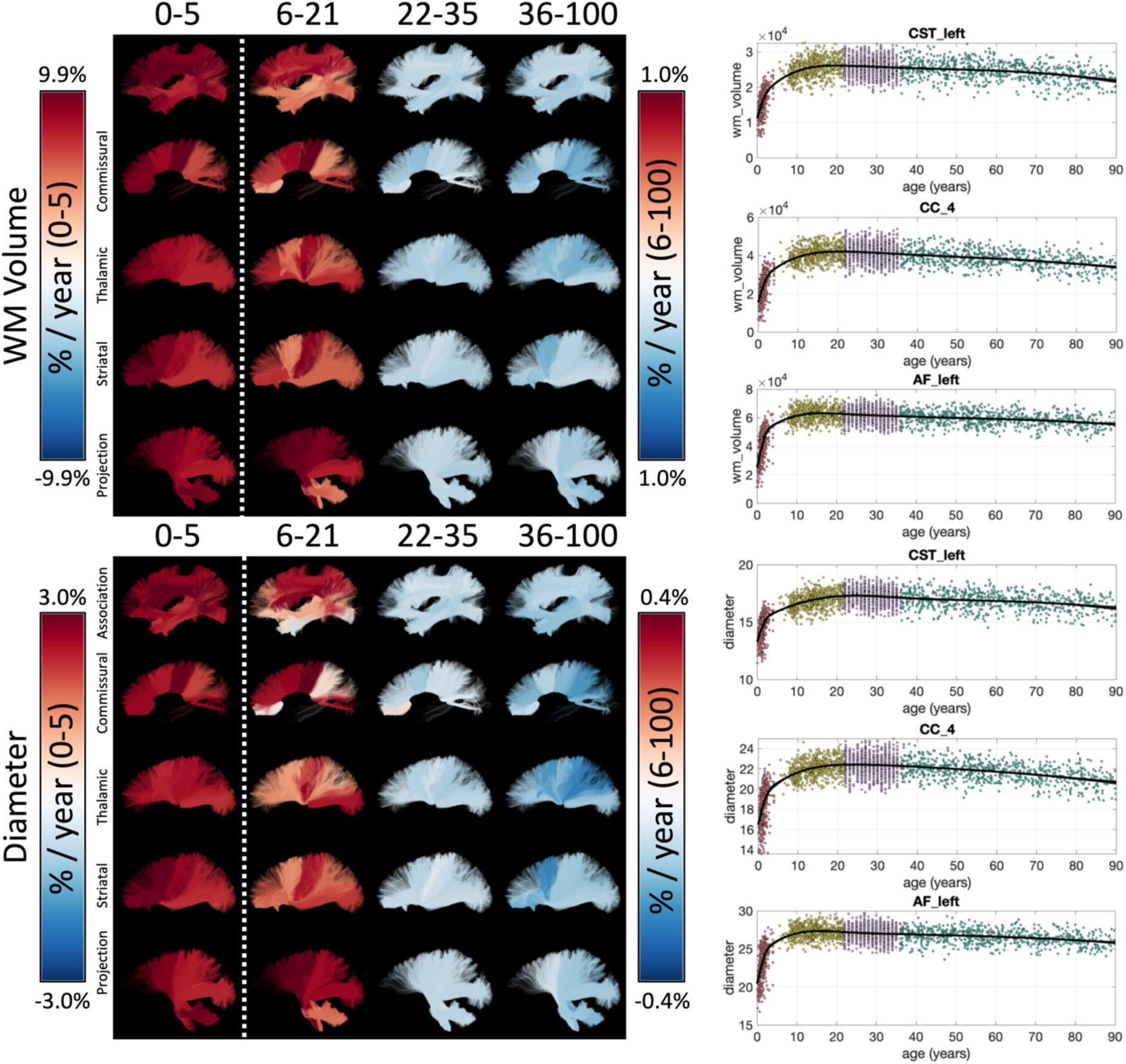
*White matter macrostructure shows unique lifespan trajectories. Lifespan trajectories of WM Volume (top) and bundle diameter (bottom) are shown colored based on the % change per year for each cohort (left) and data points are plotted against age for three selected pathways (right). Note 0-5 cohort is displayed on a different color scale than other cohorts*.

Trajectories of additional macrostructural features are given in **Supplementary Figure 3**. The % difference per year of all white matter macrostructural features is shown in **Figure 5**, for all bundles, and all cohorts. Most features of volumes, length, and surface area increase rapidly during Infancy and continue in adolescence although with reduced magnitudes. Notably, many pathways with connections to the gyri within the frontal lobe, in particular pre-and post-central cortex, indicate continued volumetric and surface area expansion into adolescence. One notable exception to the positive associations with age is elongation (length to diameter ratio), suggesting that the pathway width is increasing greater than length. A reversal of trends is clear in Young Adulthood and Aging, with nearly all features of size and shape decreasing in these cohorts, with surface area generally showing the greatest negative associations with age. In Aging, average length does not always decrease, rather decreases are associated with overall volume occupied by the pathway.

**Figure 5.**
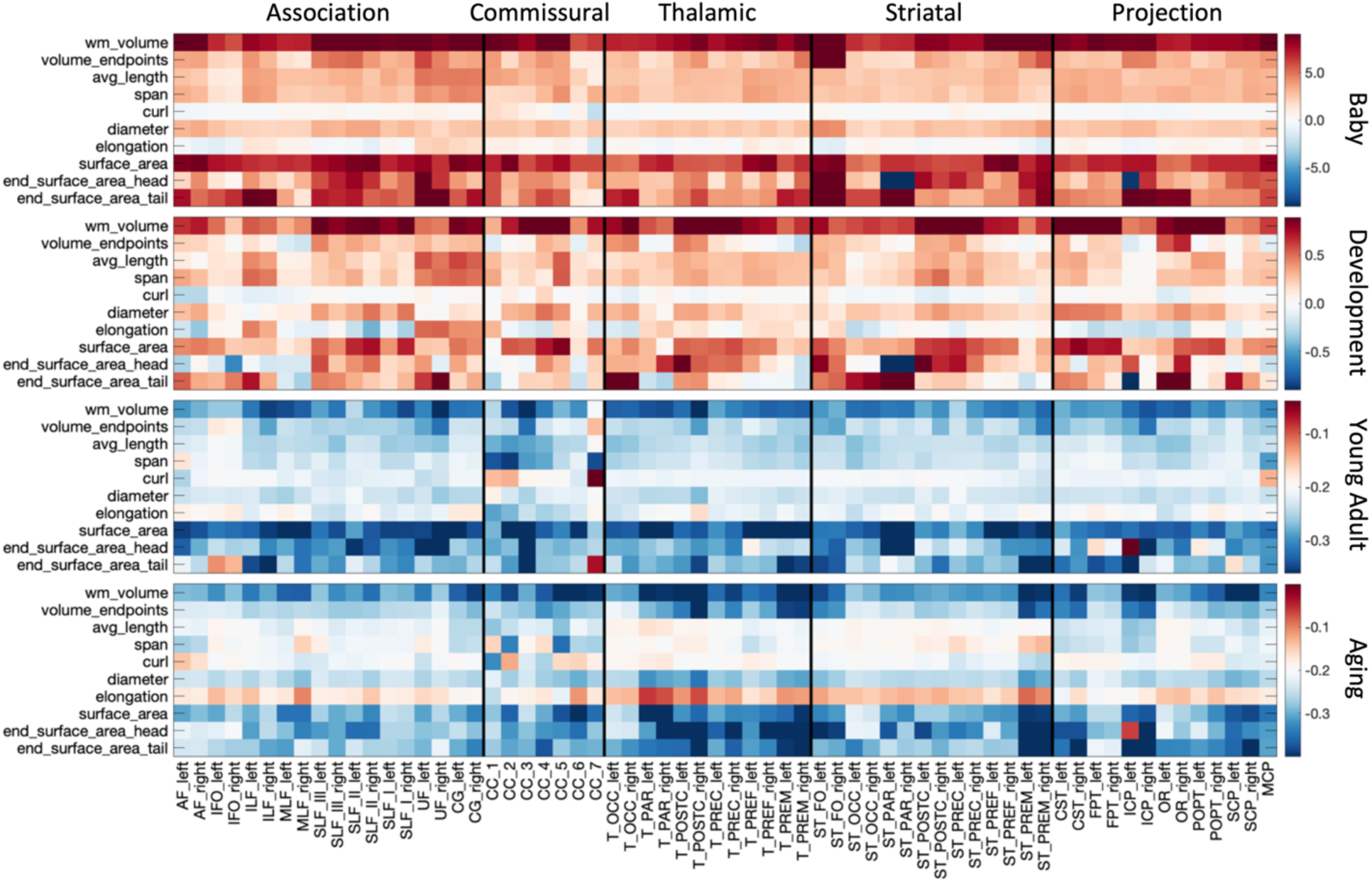
*White matter macrostructure shows different associations with age across pathways and across different lifespan stages. The % change per year of all white matter macrostructure features, for all bundles is shown for each of the four cohorts. Note, each cohort is visualized on a different scale to highlight trends across features*.

#### Cortical Structure

**Figure 6** shows the lifespan trajectories of the cortical thickness associated with each white matter bundle. Additionally, we visualize the cortical thickness of each bundle and the % difference per year for each cohort. First, different bundles have different cortical thicknesses (i.e., they connect areas with different cortical thicknesses). There is a clear pattern, with bundles connecting inferior temporal gyri, inferior frontal gyri, and superior middle frontal gyri displaying higher cortical thickness, and bundles connecting pre-and post-central gyri displaying lower cortical thickness – both in agreement with expected patterns across the cortex [17, 19]. Second, in our dataset (and with our CRCS fits) the cortical thickness decreases continually throughout the lifespan, with greater decreases in the Infant, Development, and finally the Aging datasets. Third, the difference per year varies across the pathways, with pathways connecting pre-and post-central gyri remaining relatively stable across all stages of the lifespan. Trajectories of additional cortical features are given in **Supplementary Figure 4**.

**Figure 6.**
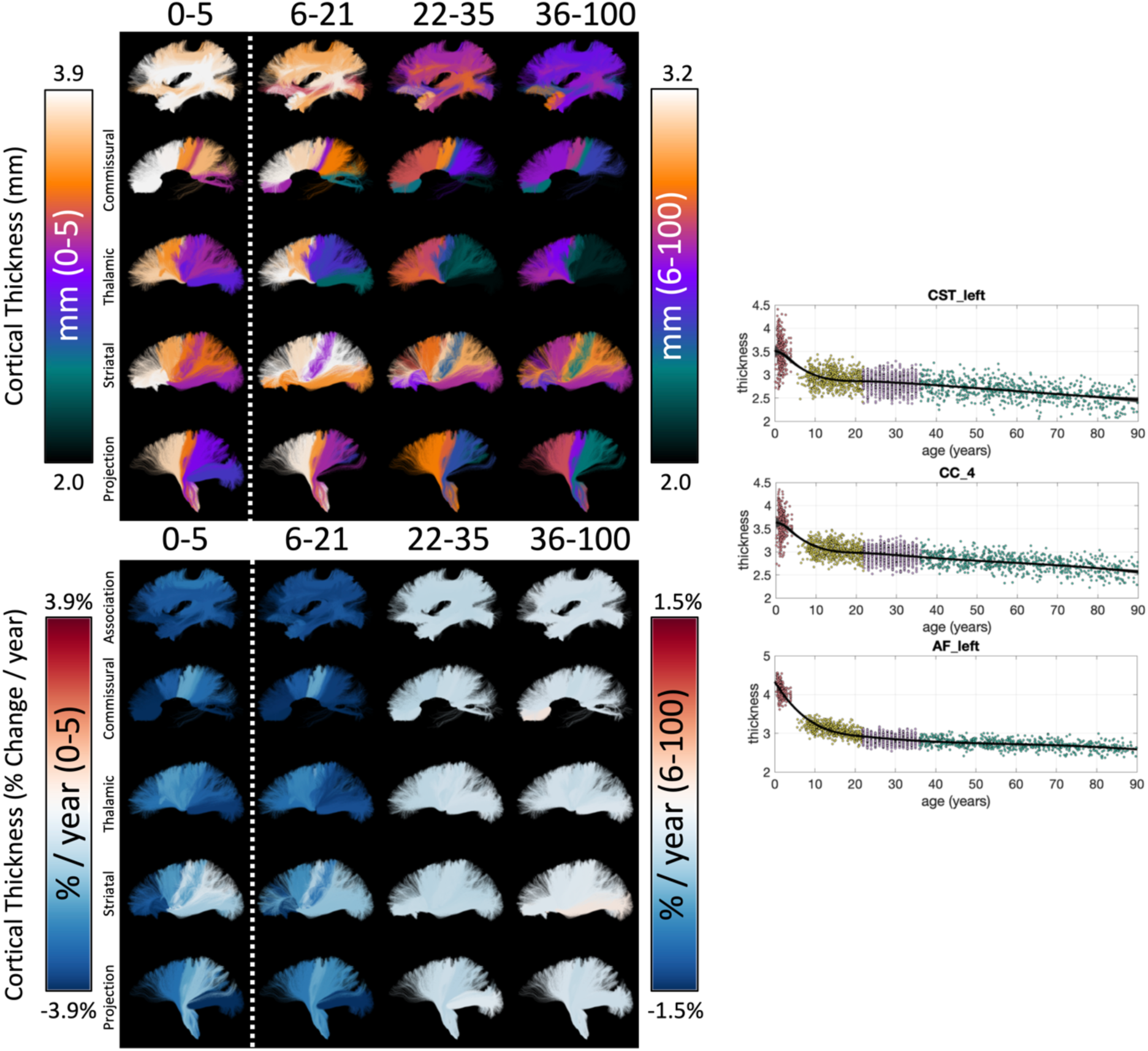
*The cortical structure associated with white matter pathways shows unique lifespan trajectories. The cortical thickness of each white matter bundle at different lifespan stages is shown (top) along with the % change per year for each cohort (bottom), and data points are plotted against age for three selected pathways (right). Note 0-5 cohort is displayed on a different color scale than other cohorts, and Cortical Thickness (in mm) is displayed as a different colormap (as opposed to the divergent colormap for % change per year)*.

The % difference per year of all cortical features associated with each pathway is shown in **Figure 7**. The Infant cohort shows large increases in cortical area and decreases in curvature and thickness for most bundles. Similarly, trends are seen in the Development cohort, with the addition of decreases in cortical volume (note again that Jacobian of White Matter and Volume are not derived for the Infant cohort). The decreases in curvature, sulcal depth, thickness, and volume continue into Young Adulthood, whereas the cortical area largely plateaus for most bundles. These trends again continue into Aging.

**Figure 7.**
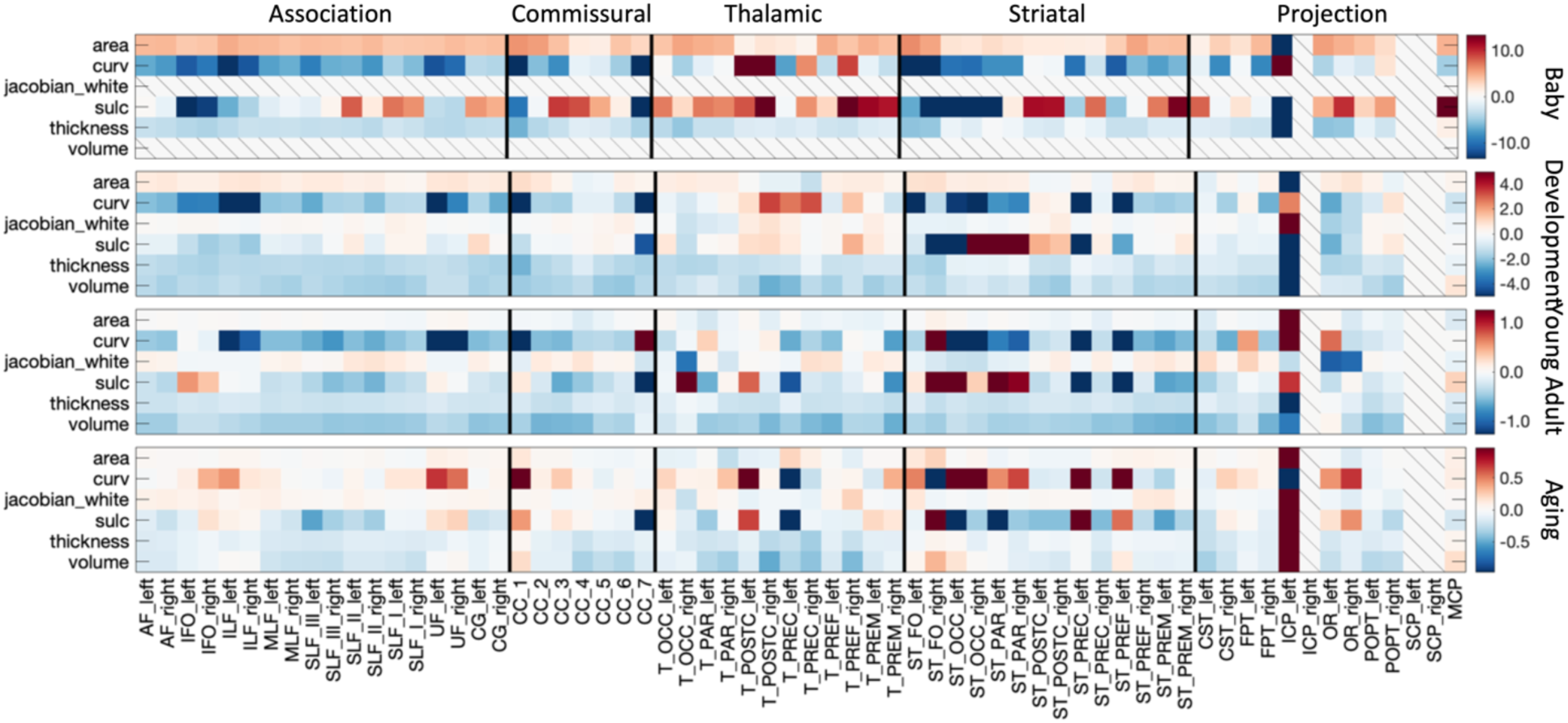
*White matter cortical structure shows different associations with age across pathways and across different lifespan stages. The % change per year of all white matter cortical features, for all bundles is shown for each of the four cohorts. Note, each cohort is visualized on a different scale to highlight trends across features, Volume and Jacobian are not given for Infant dataset, and several mid-brain pathways do not have associated cortex*.

#### Spatial gradients of age-associations

Beyond pathway-type differences, there is evidence for spatial gradients of age-associations across the brain. Here, we sorted pathways from anterior-to-posterior based on their average location in MNI-space and show selected results in **Figure 8**. Features of microstructure, macrostructure, and cortex all show significant trends that vary based on location. For example, ICVF generally shows greater associations with age for more anterior pathways, for Infant, Development, and Young Adult cohorts, with a nearly reversed trend of greater decline in Aging for the most anterior and most posterior pathways. Similarly, more anterior pathways show greater cross-sectional increases in volume during infancy, whereas the more centralized pathways show greater increases in childhood. Finally, cortical thickness shows quadratic trends with anterior-posterior position at all stages of the lifespan, largely driven by the unique trajectories of pre-and post-central gyri and supplementary motor areas which are more centralized in the anterior-to-posterior direction.

**Figure 8.**
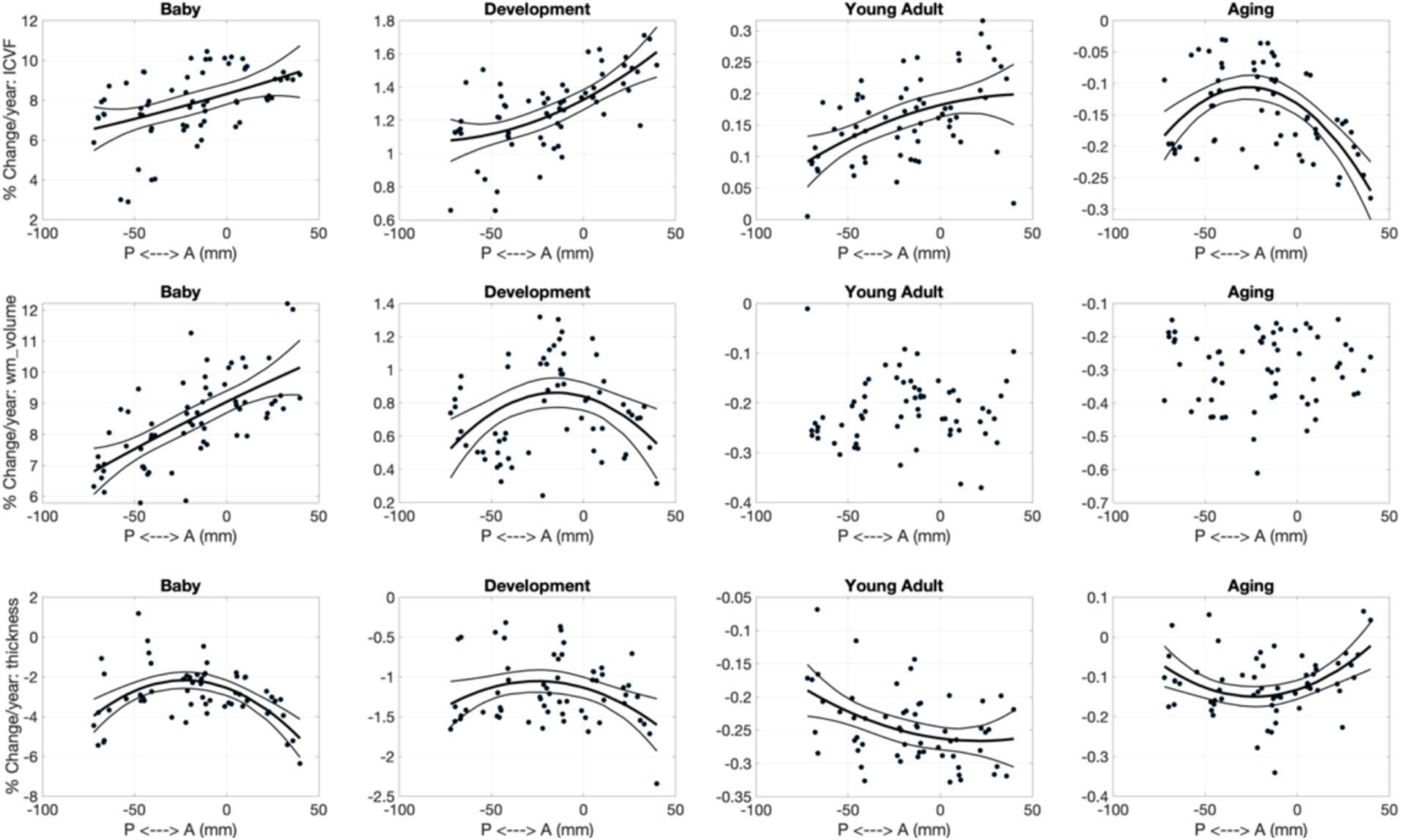
*Spatial gradients across the brain in development and degeneration patterns exist for microstructure (ICVF; top), macrostructure (WM volume; middle), and cortical structure (thickness, bottom). Linear and quadratic trends in % change per year are shown across the anterior-to-posterior direction*.

#### Relative timing of feature peak/minimum

Many features display the expected U-shaped (or inverted U-shaped) lifespan trajectory, with a peak/minimum typically occurring during young adulthood. **Figure 9** shows the relative timing of white matter feature changes for selected features, indicating when the peak or minimum is reached for each pathway, along with the 95% confidence interval based on bootstrap fits. Peak anisotropy is typically reached between 20-30 years old for association pathways, just before 20 years old for commissural pathways, and with a wide variation across others – for example many striatal, thalamic, and projection pathways show evidence of continuously positive age association of FA throughout the lifespan. ICVF and RD reverse trends (ICVF peaks, RD reaches minimum) at a later age, typically at around 40 years for most association pathways, 40-60 years for several thalamic, projection, and striatal pathways, and earlier for commissural pathways (∼20 for RD, ∼30 for ICVF). Pathway volume is much more homogenous, with volume of most bundles peaking within the early 20’s. Finally, cortical area associated with different pathways typically peaks before 20 years of age, although again some bundles (and associated cortical areas) do not show evidence for a single well-defined maximum throughout the lifespan.

**Figure 9.**
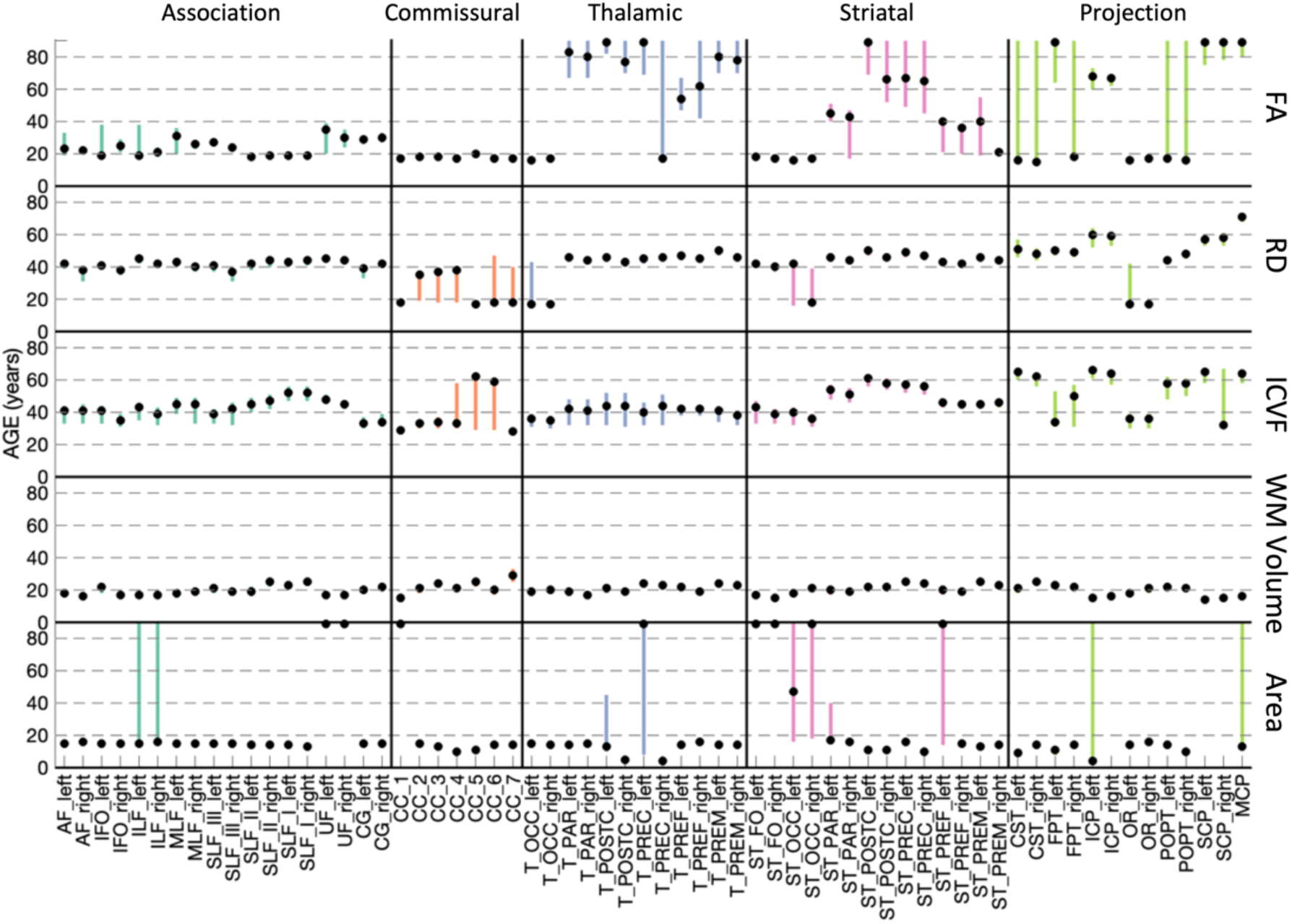
*Relative timing of white matter feature changes (i.e., peaks/minimum) for all pathways. Marker indicates age that each pathway reaches a peak/minimum. Bars show 95% bounds of peak/minimum from bootstrapping procedures. Pathways are grouped from left to right by association, commissural, thalamic, striatal, and projection/limbic pathways*.

### How are microstructural, macrostructural, and cortical features related across the lifespan?

To investigate the relationship between microstructural, macrostructural, and cortical features and how these differ across the lifespan, we calculated the cross-sectional correlation of all features to each other across a population. Results for 3 selected pathways are shown in **Figure 10**. A number of observations are clear from this figure. First, regardless of cohort, microstructure measures are strongly correlated to others – diffusivities show strong positive correlations with each other (MD, AD, RD) and strong negative correlations with FA, while ICVF shows strong positive correlations with FA. Second, again intuitively, measures of volume and area generally show strong positive correlations with each other. Third, cortical thickness shows strong positive correlations with cortical volumes and negative correlations with cortical areas, sulcal depths, and curvature. Fourth, are the more interesting relationships between different feature types. Some examples include the strong positive correlations between FA and features of white matter volumes, lengths, and areas, across all cohorts, or the positive correlations between ICVF and the same features. Fifth, different pathways do not always show the same relationships among features. For example, the relationship between cortical thickness and microstructure does not hold true for all pathways; or the unique observation that the end beginning (head) of a bundle does not always positively correlate with features at the end (tail) of the bundle (see AF_left for an example with negative correlation in Development and Young Adulthood).

**Figure 10.**
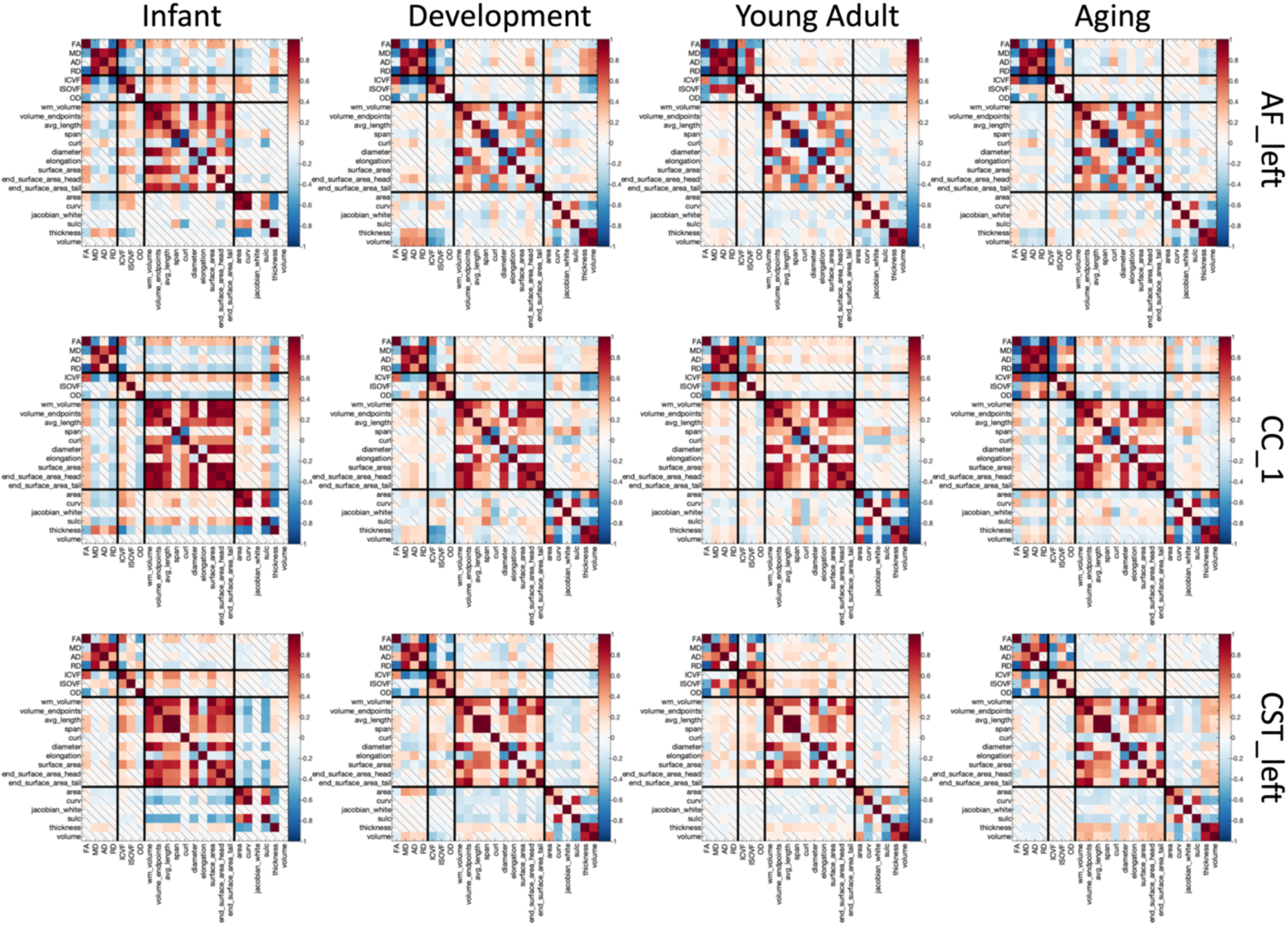
*Correlations exist among white matter features of microstructure, macrostructure, and cortex, that change with age and vary across pathways. Correlation coefficients are shown between all features for three selected pathways (top to bottom) and for each of four cohorts (left to right). Non-statistically significant correlation coefficients indicated by diagonal bar*.

Finally, the relationship between features differs at different stages of the lifespan, as seen in **Figure 10**. For example, cortical thickness (highlighted with an arrow in the figure) shows negative associations with FA and positive with diffusivities (MD, AD, RD) during Infancy and Development, but reverses relationships within the aging cohort. Feature relationships averaged across all pathways are shown in **Supplementary Figure 6**, and results for partial correlations in **Supplementary Figure 7**. All observations hold true, although with diminished magnitude for the partial correlations.

### Are age associations of different features related to each other?

We first ask, “within a cohort, do pathways with greater/smaller age-associations in one feature correspond to greater/smaller age-associations in another feature?”. The correlation coefficient of the difference-per-year (cross-sectional rates of change) of features to each other are shown in **Figure 11**, with notable observations shown as additional plots. Differences in features are strongly related to each other. During infancy, pathways with greater age-associations in white matter volume also show greater age-associations in ICVF and cortical area, pathways with greater cross-sectional decreases in MD also show greater increases in fiber diameter, and pathways with greater cortical thickness increases also experience greater RD increases. These patterns also change with age. During Aging, pathways with greater white matter volume decreases are accompanied by greater increases in ISOVF and greater decreases in cortical volume, pathways with greater decreases in ICVF experience an increase in cortical area. Uniquely, changes in the surface areas at the beginning and end of pathways are not strongly correlated across pathways.

**Figure 11.**
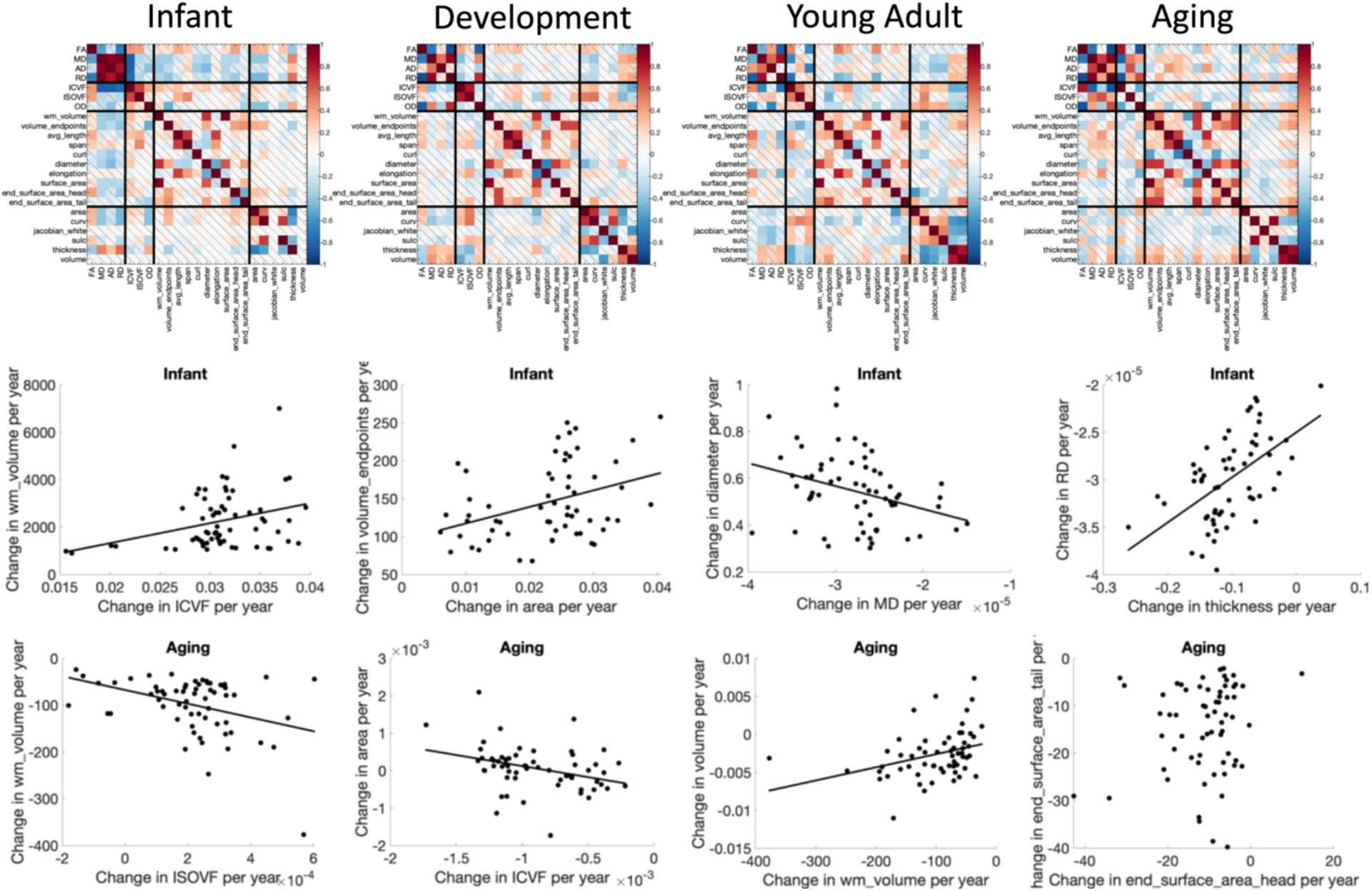
*Rates of change (association with age) of microstructure, macrostructure, and cortical features of pathways show strong relationships. Matrices show the correlation coefficients between the % change per year of features, averaged across pathways (top) and selected examples in the Infant and Aging cohorts showing the relationships between features*.

### Does white matter pathway development influence brain degeneration later in life?

Finally, we ask “do pathways with features that develop faster correspond to those that decline faster/slower in aging?”. **Figure 12** shows several examples of features where difference per year in the Development cohort are plotted against differences per year in the Aging cohort (see **Supplementary Figure 8** for all features investigated). For microstructure – pathways that show greatest age-associations in development tend to also show the greatest opposite age-associations in aging for most features. The only microstructural features that do not show statistically significant relationships between childhood and aging are AD and OD. Macrostructural features of white matter volume and surface area show similar trends, for example pathways that show the greatest age-related differences in childhood also show them in aging (i.e., CC segments, FPT, POPT, ST_PREF, and T_PREF). Finally, pathways with cortical volumes that decreases greatest in development also tend to decrease greatest in aging, for example occipital connections (OR, T_OCC, and ST_OCC), and other striatal and thalamic pathways (T_PAR, T_PREC, ST_PREC, and ST_PAR).

**Figure 12.**
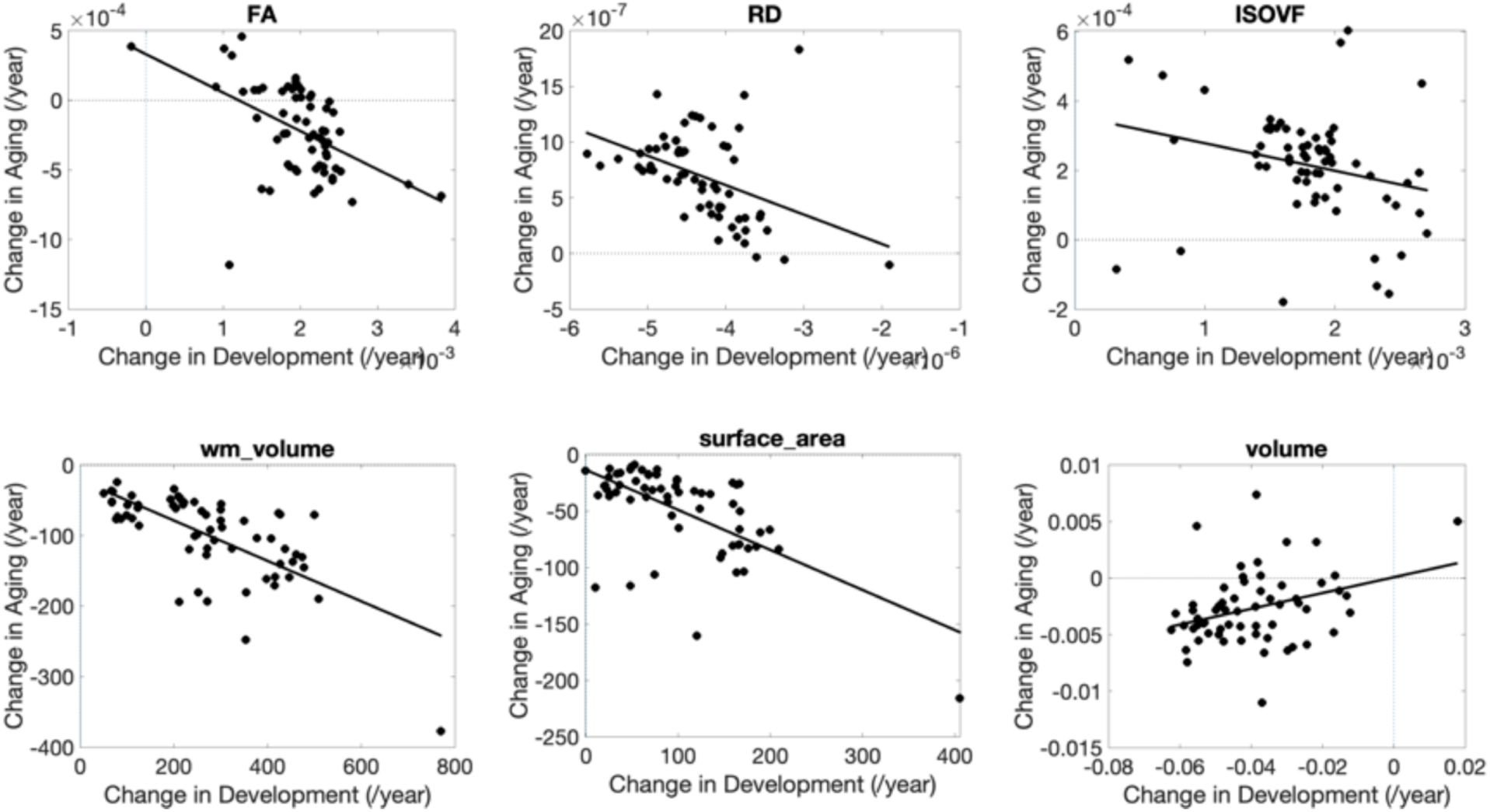
*Rates of change during development are strongly related to rates of change in aging. Selected examples showing strong linear correlations are plotted, where each datapoint represents a different pathway*.

## Discussion

We have provided a comprehensive characterization of microstructural, macrostructural, and associated cortical features of white matter bundles across the lifespan in a large cohort of normal subjects. We find that all features show unique lifespan trajectories, with rates and timing of development and degradation that vary across pathways. Certain features tend to change together, and pathway-specific trends during development are strongly related to age-related changes. Characterizing the relationships among and between different features in this way may help elucidate biological changes occurring during different stages of the lifespan and highlight atypical trajectories.

### White matter features throughout the lifespan

#### Microstructure

All *microstructure* indices demonstrated spatiotemporally varying rates of development and aging (**Figure 2**). Our results are consistent with previous DTI and NODDI-based literature, but importantly link prior studies on isolated age ranges and include more white matter pathways. Throughout infancy and childhood, FA increases and diffusivities (MD, AD, RD) decrease [3, 5], reflecting biological changes such as increased myelination and axonal packing [42], with greater rates of change observed in frontal regions and limbic connections of the brain [3, 43]. These changes continue into adulthood with similar regional patterns [4, 44]. In healthy aging, this pattern inverts, with decreases in FA and increase in diffusivities throughout much of the white matter [45-48], and more pronounced changes in frontal regions [7, 49-51]. These changes likely reflect degradation of white matter microstructure – including demyelination, disruption of axonal structure/coherence, and increased water content. RD generally shows greater relative changes than AD in infancy and development (decreasing diffusivities) as well as aging (increasing diffusivities), driving the FA changes. Biologically, this suggests that myelination and/or packing density (increases in development, decreases in aging) may be driving these changes (i.e., RD changes [52]) relative to differences in axonal structure and orientation coherence (i.e., AD changes [52]), especially with conflicting changes in the more specific OD during aging.

Rates of change across white matter pathways are not fully described by a simple anterior-to-posterior gradient nor by grouping connections (e.g., association, projection, or commissural pathways). For example, commissural, striatal, and thalamic pathways had stronger associations with age in frontal pathways during development and aging, whereas spatial trends were not clearly visible across association and projection pathways. Further, regional variation is more pronounced in aging. For example, while FA decreases during aging in many pathways, some pathways remain relatively stable or even increase in old age, particularly pathways to the pre-and post-central gyri (thalamic, striatal, and corticospinal tracts) and those associated with the cerebral peduncles. These same pathways typically show decreasing OD - reflecting either an increased coherence of neurites, or selective degeneration of cross fibers in non-projection pathways [53] – and decrease in overall volume fraction.

Patterns of change varied with age, and there is approximately a factor of ∼5 decrease in magnitude in percent change per year from infants to children, and a similar factor of ∼5 decrease from children to aging. This is broadly consistent with prior studies in more isolated age ranges [5, 12, 14, 54], but here we are able to combine multiple cohorts and demonstrate patterns across the entire lifespan.

#### Macrostructure

All studied association, projection, thalamic, striatal, and commissural pathways showed positive associations of volume with age in infancy and childhood, followed by atrophy beginning during young adulthood that continued (often nonlinearly) during aging. These findings agree with the literature which shows initially large increases in volume for most tracts during development [4], with several pathways (CC, ILF, CST) continuing post-adolescent growth into adulthood [4, 8], followed by atrophy in aging [8, 47].

Other macrostructural properties of pathways, including length, area, volume of full bundles, endpoints, and/or the trunk of bundles [16], have not been thoroughly investigated, despite their importance for fully understanding brain growth and aging. Our main finding for these other metrics, similar to volume, is that trajectories vary across pathways of the brain (**Figure 4**). The volume of endpoints shows similar trends to overall volume (as above), although of smaller magnitude, meaning that the volume of the trunk decreases faster than that of endpoints. Length shows strong positive associations with age during childhood, negative associations during young adulthood, and generally negative associations during aging, but some small but positive associations were apparent for striatal and thalamic pathways near the expanding ventricles. Surface area at the beginning of bundles did not always show similar age-associations to surface areas at the end of bundles, which means that the cortical surface areas that these connect do not atrophy at similar rates, even though they share similar connections. This might suggest a spatial gradient driving atrophy that is not driven by structural connections. Similarly, structural covariance [55] across the cortex is also not exclusively driven by connectivity alone [56].

#### Cortical features

Here, for the first time, we directly associate specific association, projection, and commissural fiber pathways of the brain with the cortical regions that they connect, assigning cortical feature measurements to each bundle. The cortical thickness varied dramatically across bundles (**Figure 6**). In general, association, commissural, and striatal pathways connecting frontal/prefrontal regions were associated with the greatest thicknesses, while those connecting with visual and motor cortices are associated with the lowest thicknesses. Second, cortical thickness of bundles decreased continually across the lifespan, with rates of change varying across pathways. This, intuitively, matches expected trends of cortical regions, with large decreases in infancy with a steeper slope up to the third decade of life and a gradual decrease thereafter [19].

Other features of the cortex similarly offer unique insight into white matter development and interaction with the cortex. For example, much like the white matter surface area increases in development and decreases thereafter, so does the expansion and contraction of the cortical surface area, although the changes have largely plateaued in the aging cohort [57]. Measures of cortical volume associated with white matter pathways similarly show large decreases in development (while the white matter volume increases), plateaus in young adulthood, and a continual decrease in aging (again as expected based on cortical studies [57, 58]). Measures of curvature and sulcal depth are less intuitive. For example, most pathways are associated with cortex with a negative curvature value (i.e., associated with gyri; see Supplementary Figure 4), but the *change* in curvature increases (tends more towards 0; **Figure 7**) in development and levels off with age, although tends to remain overall negative in magnitude. While this could be interpreted as a biological result suggesting morphogenesis and development of pathways tends to form gyral-followed-by-sulcal connections, or that these long-range pathways primarily connect gyri [59, 60], it is likely influenced by tractography biases favoring gyral connections [61] – however, it could also be some combination of true favoring of gyral connections with age and greater bias in infancy/development datasets.

#### Gradients across the brain

Both visually (**Figures 2, 4, and 6**) and using regression (**Figure 8**), we found gradients of change across the brain for many features. These were often quadratic, with greater change observed in the most anterior and most posterior pathways. The examples in **Figure 8** include ICVF, which shows a positive association with age in anterior pathways during infancy, childhood, and young adulthood, and a quadratic trend in aging; the strongest age-associations were in the most anterior and posterior pathways. Total white matter volume had linear and quadratic trends in infancy and development, respectively, but no clear trends into and past adulthood. Cortical thickness showed quadratic trends at all developmental stages.

Previous studies have described spatial gradients in developmental and degeneration patterns. During childhood, development proceeds along the posterior-to-anterior [10, 62], inferior-to-superior [42, 63], and medial-to-lateral [64] axes of the brain. Similar results have been described in aging, with faster decline along the inferior-to-superior [65, 66] and posterior-to-anterior axes with the frontal white matter particularly vulnerable [8, 66-70]. These observations (while not universally observed [13, 46, 48, 71]) are typically explained as a function of the later developing parts of the brain – which support higher order cognitive abilities and may be more vulnerable to age-related effects than those that support sensory or motor processes. Our results extend these observations to multi-compartment diffusion indices, macrostructure, and cortical features, and further partially explain minor discrepancies in the literature.

While most studies above describe gradients using a voxel-wise approach, we have chosen to select the center of mass of pathways in a standard space – however, pathways are not organized as parallel and perfectly ordered structures, and may in fact overlap with others within the same voxels [72], and gradients as a function of position may not be fully appropriate. Regardless, these results highlight the fact that linear gradients of change may be a simplification of the development and aging process and highlight the need for pathway-specific normative trajectories across the lifespan.

#### Milestones and timings

Identifying neurodevelopmental milestones is a necessary step in characterizing normative trajectories, providing insight into biological changes in the brain, and identifying abnormal trajectories [73]. Here we show that total white matter volume plateaus much before FA, which reverses trends before RD and ICVF. The total white matter volume generally peaks around 18-25 years old, which well-matches the total white matter volume trends [74] which reverse at <25 years old. Yet, cortical surface area peaks before white matter volume. Second, all features show trends that are well-grouped by pathway type. For example, FA peaks for commissural fibers, association fibers, followed by the (heterogenous peaks of striatal fibers) and finally thalamic and projection fibers. Third, these projection fibers, as well as some striatal and thalamic connections, vary widely across microstructure and cortical features, which may reflect their relative stability over the lifespan [29, 67].

#### Feature relationships

Microstructure features were strongly related to other microstructure features. DTI is sensitive to a number of biological properties of the tissue [52], so it is not surprising that microstructural measures were strongly correlated to one another [75, 76] at all points in the lifespan. Macrostructural measures also showed strong relationships to microstructure. Because volume loss (as well as losses in lengths, areas), particularly in aging, occurs due to loss of myelinated fibers [77], it is not surprising that volume was positive associated with ICVF, negatively associated with RD/MD, and negatively associated with ODI during aging. Finally, we found relationships between both micro/macrostructure and cortical features. As the cortex thins - likely due to loss of dendritic arbors [78-80] in combination with reductions in synapses, and shrinking of soma [81] (in aging) or due to myelination (in development) [82] - the white matter also undergoes changes. For example, cortical thickness is negatively associated with MD, AD, RD, and ISOVF (reductions in white matter myelination concurrent with reduced thickness) and positive associations with white matter volume, diameter, and endpoint volume, in particular. Finally, these associations are not the same across all pathways. While general trends described above hold true, different pathways show different associations – the process of development and aging occurs in different ways for different pathways.

Comparing rates of change between features has also been performed, for example finding a strong positive relationship between diffusivity developmental change and R1 (a relaxometry measure of myelin content) developmental change [9], or a strong negative relationship between rate of cortical thickness decrease and rate of FA increase in development. These findings strengthen the hypothesis that cortical thinning is likely a result of cortical myelination (affecting the apparent white-gray matter boundary) rather than cortical pruning [82-84], showing synchronous cortical myelination and white matter microstructural enhancement. In our study, we find similar synchronous rates of change between features during all stages of the lifespan. During infancy/development it is intuitive that pathways with greater rates of change in macrostructure (bundle volume, bundle diameter) have corresponding greater changes in microstructure (ICVF, MD, respectively), or that greater decreases in cortical thickness (greater pruning) is associated with greater decreases in RD (i.e., greater rates of myelination and/or greater packing density due to axon caliber enlargement [85]). Degradation during aging shows similar intuitive trends. Pathways with greater rates of change of putative interstitial spaces or diffusivities (ISOVF) experience the greatest atrophy, and greatest rates of decrease of cortical volume. One interesting observation is that the rates of change of the beginning and end of bundles do not strongly correlate during aging, meaning that connected cortical areas do not necessarily experience changes in the cortex (i.e., myelination and/or loss of dendritic arbors) at the same rate.

#### Development and Aging relationships

For many features, a steeper slope during development corresponded to steeper changes in aging. This is consistent with prior literature relating peak age to the *rate of decline* during aging [67], in contrast to the retrogenesis principle that postulates that pathways that mature later are more susceptible to degeneration during aging [31]., This study provides evidence that there is a strong pathway-specific relationship between these two crucial stages of the lifespan, where the rates of development of white matter features influence the rates of brain degeneration later in life.

#### Limitations

A limitation of the current study is differences in acquisition parameters and sites, which can lead to different microstructure and macrostructural measures [38, 86-88]. The addition of more data, from different centers, and with overlaps in age/sex will ensure more appropriate harmonization and generalizability of findings.

## Conclusion

In conclusion, state-of-the art tractography of 63 white matter pathways over the entire lifespan (0 to 100 years of age) in a large sample demonstrated age-related changes in white matter pathways, with variation in timing and magnitude of development and aging processes. There are strong relationships between features of white matter pathways, suggesting synchronous changes in white matter microstructure, white matter macrostructure, and their connecting cortex. Finally, there are pathway-specific trends during development are strongly related to age-related decline. Together, this complete characterization of white matter provides normative data that will be useful for studying normal development and degeneration or compared against abnormal processes in disease or disorder.

## Data availability

The data used in this study come from the Human Connectome Project (Essen et al. 2012), including HCP Young Adult, HCP Aging, and HCP Development cohorts, which are freely available after appropriate data usage agreements. See the following for data download: HCP Young Adult (https://www.humanconnectome.org/study/hcp-young-adult/document/1200-subjects-data-release), HCP Aging and Development (https://nda.nih.gov/general-query.html?q=query=featured-datasets:HCP%20Aging%20and%20Development). Data derivatives from this study are packaged and ready to be shared with any HCP-authorized investigator with appropriate data usage agreement from the Human Connectome Project.

## Acknowledgements

This work was conducted in part using the resources of the Advanced Computing Center for Research and Education at Vanderbilt University, Nashville, TN. This work was supported by the National Institutes of Health (NIH) under award numbers K01EB032989 (KGS), R01EB017230 (BAL), K01AG073584 (DBA), R01MH123201 (JCG and BAL), and in part by the National Center for Research Resources, Grant UL1 RR024975–01.

## Competing Interests

The authors have no relevant financial or non-financial interests to disclose.

## Author Contributions

All authors contributed to the study conception and design. All authors commented on previous versions of the manuscript. All authors read and approved the final manuscript.

## Ethic Approval

The data used in this study come from the Human Connectome Project (Essen et al. 2012), including HCP Young Adult, HCP Aging, and HCP Development cohorts, which are freely available after appropriate data usage agreements. This study accessed only de-identified patient information, an

## Appendix

The bundles resulting from the TractSeg pipeline are given as a list below, with acronyms used in the text.

Association pathways: Arcuate fascicle left (AF_L); Arcuate fascicle right (AF_R); Cingulum left (CG_L); Cingulum right (CG_R); Inferior occipito-frontal fascicle left (IFO_L); Inferior occipito-frontal fascicle right (IFO_R); Inferior longitudinal fascicle left (ILF_L); Inferior longitudinal fascicle right (ILF_R); Middle longitudinal fascicle left (MLF_L); Middle longitudinal fascicle right (MLF_R); Superior longitudinal fascicle III left SLF_III_L); Superior longitudinal fascicle III right (SLF_III_R); Superior longitudinal fascicle II left (SLF_II_L); Superior longitudinal fascicle II right (SLF_II_R); Superior longitudinal fascicle I left (SLF_I_L); Superior longitudinal fascicle I right (SLF_I_R); Uncinate fascicle left (UF_L); Uncinate fascicle right (UF_R).

Commissural pathways: Rostrum (CC_1); Genu (CC_2); Rostral body (Premotor)(CC_3); Anterior midbody (Primary Motor) (CC_4); Posterior midbody (Primary Somatosensory) (CC_5); Isthmus (CC_6); Splenium (CC_7);

Projections/Cerebellar pathways: Corticospinal tract left (CST_L); Corticospinal tract right (CST_R); Fronto-pontine tract left (FPT_L); Fronto-pontine tract right (FPT_R); Inferior cerebellar peduncle left (ICP_L); Inferior cerebellar peduncle right (ICP_R); Middle cerebellar peduncle (MCP); Optic radiation left (OR_L); Optic radiation right (OR_R); Parietooccipital pontine left (POPT_L); Parieto-occipital pontine right (POPT_R); Superior cerebellar peduncle left (SCP_L); Superior cerebellar peduncle right (SCP_R);

Striatal pathways: Striato-fronto-orbital left (ST_FO_L); Striato-fronto-orbital right (ST_FO_R); Striato-occipital left (ST_OCC_L); Striato-occipital right (ST_OCC_R); Striato-parietal left (ST_PAR_L); Striato-parietal right (ST_PAR_R); Striatopostcentral left (ST_POSTC_L); Striato-postcentral right (ST_POSTC_R); Striato-precentral left (ST_PREC_L); Striato-precentral right (ST_PREC_R); Striato-prefrontal left (ST_PREF_L); Striato-prefrontal right (ST_PREF_R); Striato-premotor left (ST_PREM_L); Striato-premotor right (ST_PREM_R); Thalamo-occipital left (T_OCC_L);

Thalamic pathways: Thalamo-occipital right (T_OCC_R); Thalamo-parietal left (T_PAR_L); Thalamo-parietal right (T_PAR_R); Thalamopostcentral left (T_POSTC_L); Thalamo-postcentral right (T_POSTC_R); Thalamo-precentral left (T_PREC_L); Thalamo-precentral right (T_PREC_R); Thalamo-prefrontal left (T_PREF_L); Thalamo-prefrontal right (T_PREF_R); Thalamo-premotor left (T_PREM_L); Thalamo-premotor right (T_PREM_R);

## Notes

### Competing Interest Statement

The authors have declared no competing interest.

## References

1. Pierpaoli, C., et al., Diffusion tensor MR imaging of the human brain. Radiology, 1996. 201(3): p. 637–48.

2. Basser, P.J., J. Mattiello, and D. LeBihan, MR diffusion tensor spectroscopy and imaging. Biophys J, 1994. 66(1): p. 259–67.

3. Reynolds, J.E., et al., Global and regional white matter development in early childhood. Neuroimage, 2019. 196: p. 49–58.

4. Lebel, C. and C. Beaulieu, Longitudinal development of human brain wiring continues from childhood into adulthood. J Neurosci, 2011. 31(30): p. 10937–47.

5. Cancelliere, A., et al., DTI values in key white matter tracts from infancy through adolescence. AJNR Am J Neuroradiol, 2013. 34(7): p. 1443–9.

6. Fjell, A.M., et al., The relationship between diffusion tensor imaging and volumetry as measures of white matter properties. Neuroimage, 2008. 42(4): p. 1654–68.

7. Isaac Tseng, W.Y., et al., Microstructural differences in white matter tracts across middle to late adulthood: a diffusion MRI study on 7167 UK Biobank participants. Neurobiol Aging, 2021. 98: p. 160–172.

8. Lebel, C., et al., Diffusion tensor imaging of white matter tract evolution over the lifespan. Neuroimage, 2012. 60(1): p. 340–52.

9. Yeatman, J.D., B.A. Wandell, and A.A. Mezer, Lifespan maturation and degeneration of human brain white matter. Nat Commun, 2014. 5: p. 4932.

10. Westlye, L.T., et al., Life-span changes of the human brain white matter: diffusion tensor imaging (DTI) and volumetry. Cereb Cortex, 2010. 20(9): p. 2055–68.

11. Zhang, H., et al., NODDI: practical in vivo neurite orientation dispersion and density imaging of the human brain. Neuroimage, 2012. 61(4): p. 1000–16.

12. Zhao, X., et al., Brain Development From Newborn to Adolescence: Evaluation by Neurite Orientation Dispersion and Density Imaging. Front Hum Neurosci, 2021. 15: p. 616132.

13. Mah, A., B. Geeraert, and C. Lebel, Detailing neuroanatomical development in late childhood and early adolescence using NODDI. PLoS One, 2017. 12(8): p. e0182340.

14. Cox, S.R., et al., Ageing and brain white matter structure in 3,513 UK Biobank participants. Nat Commun, 2016. 7: p. 13629.

15. Beck, D., et al., White matter microstructure across the adult lifespan: A mixed longitudinal and cross-sectional study using advanced diffusion models and brain-age prediction. Neuroimage, 2021. 224: p. 117441.

16. Yeh, F.C., Shape analysis of the human association pathways. Neuroimage, 2020. 223: p. 117329.

17. Fischl, B. and A.M. Dale, Measuring the thickness of the human cerebral cortex from magnetic resonance images. Proc Natl Acad Sci U S A, 2000. 97(20): p. 11050–5.

18. Brickman, A.M., et al., Structural MRI covariance patterns associated with normal aging and neuropsychological functioning. Neurobiol Aging, 2007. 28(2): p. 284–95.

19. Frangou, S., et al., Cortical thickness across the lifespan: Data from 17,075 healthy individuals aged 3-90 years. Hum Brain Mapp, 2022. 43(1): p. 431–451.

20. Roe, J.M., et al., Tracing the development and lifespan change of population-level structural asymmetry in the cerebral cortex. Elife, 2023. 12.

21. Wang, F., et al., Developmental topography of cortical thickness during infancy. Proc Natl Acad Sci U S A, 2019. 116(32): p. 15855–15860.

22. Williams, L.Z.J., et al., Structural and functional asymmetry of the neonatal cerebral cortex. Nat Hum Behav, 2023. 7(6): p. 942–955.

23. Fjell, A.M., et al., Development and aging of cortical thickness correspond to genetic organization patterns. Proc Natl Acad Sci U S A, 2015. 112(50): p. 15462–7.

24. Walhovd, K.B., et al., Neurodevelopmental origins of lifespan changes in brain and cognition. Proc Natl Acad Sci U S A, 2016. 113(33): p. 9357–62.

25. Di Biase, M.A., et al., Mapping human brain charts cross-sectionally and longitudinally. Proc Natl Acad Sci U S A, 2023. 120(20): p. e2216798120.

26. Wang, L., et al., Alterations in cortical thickness and white matter integrity in mild cognitive impairment measured by whole-brain cortical thickness mapping and diffusion tensor imaging. AJNR Am J Neuroradiol, 2009. 30(5): p. 893–9.

27. Cafiero, R., et al., The Concurrence of Cortical Surface Area Expansion and White Matter Myelination in Human Brain Development. Cereb Cortex, 2019. 29(2): p. 827–837.

28. Jeon, T., et al., Synchronous Changes of Cortical Thickness and Corresponding White Matter Microstructure During Brain Development Accessed by Diffusion MRI Tractography from Parcellated Cortex. Front Neuroanat, 2015. 9: p. 158.

29. Tamnes, C.K., et al., Brain maturation in adolescence and young adulthood: regional age-related changes in cortical thickness and white matter volume and microstructure. Cereb Cortex, 2010. 20(3): p. 534–48.

30. Reisberg, B., et al., Evidence and mechanisms of retrogenesis in Alzheimer’s and other dementias: management and treatment import. Am J Alzheimers Dis Other Demen, 2002. 17(4): p. 202–12.

31. Brickman, A.M., et al., Testing the white matter retrogenesis hypothesis of cognitive aging. Neurobiol Aging, 2012. 33(8): p. 1699–715.

32. Howell, B.R., et al., The UNC/UMN Baby Connectome Project (BCP): An overview of the study design and protocol development. Neuroimage, 2019. 185: p. 891–905.

33. Fischl, B., FreeSurfer. Neuroimage, 2012. 62(2): p. 774–81.

34. Zollei, L., et al., Infant FreeSurfer: An automated segmentation and surface extraction pipeline for T1-weighted neuroimaging data of infants 0-2 years. Neuroimage, 2020. 218: p. 116946.

35. Glasser, M.F., et al., The minimal preprocessing pipelines for the Human Connectome Project. Neuroimage, 2013. 80: p. 105–24.

36. Van Essen, D.C., et al., The Human Connectome Project: a data acquisition perspective. Neuroimage, 2012. 62(4): p. 2222–31.

37. Wasserthal, J., P. Neher, and K.H. Maier-Hein, TractSeg - Fast and accurate white matter tract segmentation. Neuroimage, 2018. 183: p. 239–253.

38. Fortin, J.P., et al., Harmonization of multi-site diffusion tensor imaging data. Neuroimage, 2017. 161: p. 149–170.

39. Huo, Y., et al., Mapping Lifetime Brain Volumetry with Covariate-Adjusted Restricted Cubic Spline Regression from Cross-sectional Multi-site MRI. Med Image Comput Comput Assist Interv, 2016. 9900: p. 81–88.

40. Hedman, A.M., et al., Human brain changes across the life span: a review of 56 longitudinal magnetic resonance imaging studies. Hum Brain Mapp, 2012. 33(8): p. 1987–2002.

41. Stone, C.J., [Generalized Additive Models]: Comment. Statistical Science, 1986. 1(3): p. 312–314, 3.

42. Qiu, A., S. Mori, and M.I. Miller, Diffusion tensor imaging for understanding brain development in early life. Annu Rev Psychol, 2015. 66: p. 853–76.

43. Krogsrud, S.K., et al., Changes in white matter microstructure in the developing brain--A longitudinal diffusion tensor imaging study of children from 4 to 11years of age. Neuroimage, 2016. 124(Pt A): p. 473–486.

44. Giorgio, A., et al., Age-related changes in grey and white matter structure throughout adulthood. Neuroimage, 2010. 51(3): p. 943–51.

45. Salat, D.H., et al., Age-related alterations in white matter microstructure measured by diffusion tensor imaging. Neurobiol Aging, 2005. 26(8): p. 1215–27.

46. Bennett, I.J., et al., Age-related differences in multiple measures of white matter integrity: A diffusion tensor imaging study of healthy aging. Hum Brain Mapp, 2010. 31(3): p. 378–90.

47. de Groot, M., et al., Tract-specific white matter degeneration in aging: the Rotterdam Study. Alzheimers Dement, 2015. 11(3): p. 321–30.

48. Barrick, T.R., et al., White matter structural decline in normal ageing: a prospective longitudinal study using tract-based spatial statistics. Neuroimage, 2010. 51(2): p. 565–77.

49. Schilling, K., et al., Aging and white matter microstructure and macrostructure: a longitudinal multi-site diffusion MRI study of 1,184 participants. bioRxiv, 2022: p. 2022.02.10.479977.

50. Davis, S.W., et al., Assessing the effects of age on long white matter tracts using diffusion tensor tractography. Neuroimage, 2009. 46(2): p. 530–41.

51. Sullivan, E.V., T. Rohlfing, and A. Pfefferbaum, Quantitative fiber tracking of lateral and interhemispheric white matter systems in normal aging: relations to timed performance. Neurobiol Aging, 2010. 31(3): p. 464–81.

52. Beaulieu, C., The basis of anisotropic water diffusion in the nervous system - a technical review. NMR Biomed, 2002. 15(7-8): p. 435–55.

53. Han, A., et al., Fiber-specific age-related differences in the white matter of healthy adults uncovered by fixel-based analysis. Neurobiol Aging, 2023. 130: p. 22–29.

54. Lawrence, K.E., et al., Age and sex effects on advanced white matter microstructure measures in 15,628 older adults: A UK biobank study. Brain Imaging Behav, 2021. 15(6): p. 2813–2823.

55. Mechelli, A., et al., Structural covariance in the human cortex. J Neurosci, 2005. 25(36): p. 8303–10.

56. Gong, G., et al., Convergence and divergence of thickness correlations with diffusion connections across the human cerebral cortex. Neuroimage, 2012. 59(2): p. 1239–48.

57. Storsve, A.B., et al., Differential longitudinal changes in cortical thickness, surface area and volume across the adult life span: regions of accelerating and decelerating change. J Neurosci, 2014. 34(25): p. 8488–98.

58. Sele, S., et al., Age-related decline in the brain: a longitudinal study on inter-individual variability of cortical thickness, area, volume, and cognition. Neuroimage, 2021. 240: p. 118370.

59. Nie, J., et al., Axonal fiber terminations concentrate on gyri. Cereb Cortex, 2012. 22(12): p. 2831–9.

60. Chen, H., et al., Coevolution of gyral folding and structural connection patterns in primate brains. Cereb Cortex, 2013. 23(5): p. 1208–17.

61. Schilling, K., et al., Confirmation of a gyral bias in diffusion MRI fiber tractography. Hum Brain Mapp, 2018. 39(3): p. 1449–1466.

62. Colby, J.B., J.D. Van Horn, and E.R. Sowell, Quantitative in vivo evidence for broad regional gradients in the timing of white matter maturation during adolescence. Neuroimage, 2011. 54(1): p. 25–31.

63. Storsve, A.B., et al., Longitudinal Changes in White Matter Tract Integrity across the Adult Lifespan and Its Relation to Cortical Thinning. PLoS One, 2016. 11(6): p. e0156770.

64. Hermoye, L., et al., Pediatric diffusion tensor imaging: normal database and observation of the white matter maturation in early childhood. Neuroimage, 2006. 29(2): p. 493–504.

65. Sexton, C.E., et al., Accelerated changes in white matter microstructure during aging: a longitudinal diffusion tensor imaging study. J Neurosci, 2014. 34(46): p. 15425–36.

66. Hoagey, D.A., et al., Joint contributions of cortical morphometry and white matter microstructure in healthy brain aging: A partial least squares correlation analysis. Hum Brain Mapp, 2019. 40(18): p. 5315–5329.

67. Slater, D.A., et al., Evolution of white matter tract microstructure across the life span. Hum Brain Mapp, 2019. 40(7): p. 2252–2268.

68. Sala, S., et al., Microstructural changes and atrophy in brain white matter tracts with aging. Neurobiol Aging, 2012. 33(3): p. 488–498 e2.

69. Henriques, R.N., R. Henson, and M.M. Correia, Unique information from common diffusion MRI models about white-matter differences across the human adult lifespan. arXiv preprint arXiv:2306.09942, 2023.

70. Billiet, T., et al., Age-related microstructural differences quantified using myelin water imaging and advanced diffusion MRI. Neurobiol Aging, 2015. 36(6): p. 2107–21.

71. Sullivan, E.V. and A. Pfefferbaum, Diffusion tensor imaging and aging. Neurosci Biobehav Rev, 2006. 30(6): p. 749–61.

72. Schilling, K.G., et al., Prevalence of white matter pathways coming into a single white matter voxel orientation: The bottleneck issue in tractography. Hum Brain Mapp, 2021.

73. Bethlehem, R.A.I., et al., Brain charts for the human lifespan. Nature, 2022. 604(7906): p. 525–533.

74. Schilling, K.G., et al., Superficial white matter across the lifespan: volume, thickness, change, and relationship with cortical features. bioRxiv, 2022: p. 2022.07.20.500818.

75. De Santis, S., et al., Why diffusion tensor MRI does well only some of the time: variance and covariance of white matter tissue microstructure attributes in the living human brain. Neuroimage, 2014. 89(100): p. 35–44.

76. Chamberland, M., et al., Dimensionality reduction of diffusion MRI measures for improved tractometry of the human brain. Neuroimage, 2019. 200: p. 89–100.

77. Xiong, Y.Y. and V. Mok, Age-related white matter changes. J Aging Res, 2011. 2011: p. 617927.

78. Scheibel, M.E., et al., Progressive dendritic changes in aging human cortex. Exp Neurol, 1975. 47(3): p. 392–403.

79. Nakamura, S., et al., Age-related changes of pyramidal cell basal dendrites in layers III and V of human motor cortex: a quantitative Golgi study. Acta Neuropathol, 1985. 65(3- 4): p. 281–4.

80. Vidal-Pineiro, D., et al., Cellular correlates of cortical thinning throughout the lifespan. Sci Rep, 2020. 10(1): p. 21803.

81. Esiri, M.M., Ageing and the brain. J Pathol, 2007. 211(2): p. 181–7.

82. Natu, V.S., et al., Apparent thinning of human visual cortex during childhood is associated with myelination. Proc Natl Acad Sci U S A, 2019. 116(41): p. 20750–20759.

83. Paquola, C., et al., Shifts in myeloarchitecture characterise adolescent development of cortical gradients. Elife, 2019. 8.

84. Walhovd, K.B., et al., Through Thick and Thin: a Need to Reconcile Contradictory Results on Trajectories in Human Cortical Development. Cereb Cortex, 2017. 27(2): p. 1472–1481.

85. Paus, T., Growth of white matter in the adolescent brain: myelin or axon? Brain Cogn, 2010. 72(1): p. 26–35.

86. Schilling, K.G., et al., Fiber tractography bundle segmentation depends on scanner effects, vendor effects, acquisition resolution, diffusion sampling scheme, diffusion sensitization, and bundle segmentation workflow. Neuroimage, 2021. 242: p. 118451.

87. Ning, L., et al., Cross-scanner and cross-protocol multi-shell diffusion MRI data harmonization: Algorithms and results. Neuroimage, 2020. 221: p. 117128.

88. Mirzaalian, H., et al., Inter-site and inter-scanner diffusion MRI data harmonization. Neuroimage, 2016. 135: p. 311–23.

